# Cortical surface architecture endophenotype and correlates of clinical diagnosis of autism spectrum disorder

**DOI:** 10.1101/573527

**Authors:** Bun Yamagata, Takashi Itahashi, Junya Fujino, Haruhisa Ohta, Osamu Takashio, Motoaki Nakamura, Nobumasa Kato, Masaru Mimura, Ryu-ichiro Hashimoto, Yuta Aoki

## Abstract

**Aim:** Prior structural MRI studies demonstrated atypical gray matter characteristics in siblings of individuals with autism spectrum disorder (ASD). However, they did not clarify which aspect of gray matter presents the endophenotype. Further, because they did not enroll siblings of TD people, they underestimated the difference between individuals with ASD and their unaffected siblings. The current study aimed to solve these questions.

**Methods:** We recruited 30 pairs of adult male siblings (15 of them have an ASD endophenotype, other 15 pairs not) and focused on four gray matter parameters: cortical volume and three surface-based parameters (cortical thickness, fractal dimension, and sulcal depth [SD]). First, we sought to identify a pattern of an ASD endophenotype, comparing the four parameters. Then, we compared individuals with ASD and their unaffected siblings in the cortical parameters to identify neural correlates for the clinical diagnosis accounting for the difference between TD siblings.

**Results:** A sparse logistic regression with a leave-one-pair-out cross-validation showed the highest accuracy for the identification of an ASD endophenotype (73.3%) with the SD compared with the other three parameters. A bootstrapping analysis accounting for the difference in the SD between TD siblings showed a significantly large difference between individuals with ASD and their unaffected siblings in six out of 68 regions-of-interest accounting for multiple comparisons.

**Conclusions:** This proof-of-concept study suggests that an ASD endophenotype emerges in SD and that neural correlates for the clinical diagnosis can be dissociated from the endophenotype when we accounted for the difference between TD siblings. (248/250 words)

## Introduction

Autism spectrum disorder (ASD) is a neurodevelopmental disorder characterized by deficits in social interaction.^1^ A number of neuroimaging studies focusing on gray matter have demonstrated atypical cortical structure among individuals with ASD.^2, 3^ Reflecting on the deficit, they demonstrated atypical features within the “social brain”.^4–6^

When examining cortical structure, surface-based cortical matrices, such as cortical thickness (CT), fractal dimension (FD; a marker of cortical complexity), and sulcal depth (SD) are of particular interest. Unlike the conventional approach to examine regional gray matter volume, these surface-based cortical measures focus on cortical folding.^7–10^ Since they reflect different aspects of cortical architecture and stem from different genetic and cellular mechanisms in the brain,^11, 12^ they have the potential to provide a more complete picture of the pathophysiology involved in the cortical architecture of ASD than gray matter volume.

ASD is a heritable condition ^13, 14^ and shows familial aggregation.^15^ While monozygotic twins are not always concordant for an ASD diagnosis, biological full siblings of individuals with ASD are 10 to 20 times more likely to develop ASD than the general population.^14, 16^ In addition, even first-degree relatives of individuals with ASD who do not satisfy the diagnostic criteria of ASD often have subclinical autistic traits.^17^ Such observations suggest that genetic ASD traits are complex, and unaffected siblings of individuals with ASD inherit the genetic traits associated with ASD.

Endophenotype is a measurable and biological characteristic that reflects the genetic liability for a disease and exists between genes and clinical phenotypes in both affected individuals and their unaffected relatives.^18^ Because of its complex genetic contribution, characterizing an ASD endophenotype is particularly important as it may provide an objective intermediate marker and insight into the pathophysiology of ASD. Since genetic factors are responsible for a significant amount of variation in brain anatomy,^19–21^ shared alterations in brain morphology between individuals with ASD and their unaffected siblings are likely to provide an informative endophenotype. Indeed, to identify an ASD endophenotype, prior neuroimaging studies enrolled individuals with ASD, their unaffected siblings, and individuals with typically developing (TD) people. ^22–24^ These studies compared three groups in MRI parameters and regarded the atypical findings shared by the individuals with ASD and their unaffected siblings as an ASD endophenotype.

Despite the importance of identifying the ASD endophenotype, previous studies have had some potential concerns. First, although some of the prior studies utilized the univariate analyses, it might be oversimplified given the complex and large brain network underlying ASD symptoms. In addition, many of the prior studies focused on only one parameter of gray matter. Second, given that people with an ASD endophenotype do not always develop a clinical ASD diagnosis, neural correlates for the clinical diagnosis of ASD might be dissociable from its endophenotype. However, the previous studies with three groups (i.e. individuals with ASD, their unaffected siblings, and TD people) did not account for the similarity between siblings that exist between TD siblings, which may result in underestimation of the difference between individuals with ASD and their unaffected siblings. To overcome these limitations, we enrolled 60 participants, consisting of 30 people with an ASD endophenotype (15 individuals with ASD and 15 of their unaffected siblings) and 30 people without (15 pairs of TD siblings).

## Methods

The aim of this study was two-fold. First, to describe the pattern of an ASD endophenotype, we classified pairs of people with an ASD endophenotype (i.e., individuals with ASD and their unaffected siblings) and pairs of people without (i.e., TD siblings) using multivariate approaches with volume-based and surface-based cortical parameters including cortical volume (CV), CT, FD, and SD. Then, we compared the performance of these parameters to understand which aspect of the cortex presents an ASD endophenotype. Second, we performed a bootstrapping analysis to dissociate the neural basis of the clinical diagnosis of ASD from the endophenotype. Practically, we examined whether the difference between individuals with ASD and their unaffected siblings was substantially large when we account for the difference between TD siblings.

### Participants

Details of the participants are available elsewhere.^25^ Briefly, a total of 60 adult males, consisting of 30 pairs of biological siblings, were included in the present study (Tables 1, 2). Thirty people had an ASD endophenotype. Specifically, 15 pairs of participants were discordant for the diagnosis of ASD: one of the siblings was affected with ASD, and the other was unaffected. Another 30 people did not have an ASD endophenotype and consisted of 15 pairs of TD siblings. None of the TD siblings had a family member who had been diagnosed as having ASD. The clinical diagnosis of ASD was based on the DSM-IV-TR.^26^ The diagnosis was further supported by the Autism Diagnostic Observation Schedule (ADOS).^27^ In addition, to confirm the absence of a diagnosis of ASD in the unaffected sibling, the parents of the siblings discordant for the diagnosis of ASD were interviewed using the Autism Diagnostic Interview-Revised (ADI-R).^28^ We utilized the Edinburgh Handedness Inventory to evaluate handedness.^29^ The IQ of each participant was assessed using either the Wechsler Adult Intelligence Scale-Third Edition or the WAIS-revised.^30, 31^ All the participants fulfilled the Japanese version of the Autism-Spectrum Quotient (AQ).^32^ Two participants with ASD had psychiatric comorbidities: one had attention-deficit/hyperactivity disorder, and another had learning disabilities. Five of the participants were taking medication at the time of scanning: three were taking benzodiazepine, three were taking anti-depressants, and one was taking a psycho-stimulant. The absence of an Axis I diagnosis per the DSM-IV-TR and a history of psychotropic medication were confirmed.^33^ The exclusion criteria for all the participants were known genetic diseases, neurological disorders, history of significant head trauma, or an estimated full IQ of 80 or below. Written informed consent was obtained from all the participants after they had received a complete description of the study. The Ethics Committee of Showa University approved the study protocol. The study was prepared in accordance with the ethical standards of the Declaration of Helsinki.

### Image acquisition and preprocessing

MR images were acquired using a 3.0-T MRI scanner (MAGNETON Verio, Siemens Medical Systems, Erlangen, Germany) with a 12-channel head coil. High-resolution T1-weighted images were acquired using an MPRAGE sequence (TR: 2.3 s, TE: 2.98 ms, flip angle: 9, FOV: 256 mm, matrix size: 256 × 256, slice thickness: 1 mm, 240 sagittal slices, voxel size: 1 × 1× 1 mm). All the images were first visually checked for scanner artifacts and anatomical anomalies. Structural MRI data were preprocessed using the CAT 12 toolbox (Computational Anatomy Toolbox 12; Structural Brain Mapping group, Jena University Hospital, Jena, Germany) implemented in SPM12 (Statistical Parametric Mapping, Institute of Neurology, London, UK) to obtain the regional volume data. All the T1-weighted images were corrected for bias-field inhomogeneities, then segmented into gray matter (i.e., CV), white matter and cerebrospinal fluid and spatially normalized using the DARTEL algorithm.^34^ The final resulting voxel size was 1.5 × 1.5 × 1.5 mm. For quality assurance, the resulting images were checked for homogeneity. Segmented and normalized data were smoothed with a Gaussian kernel of 8 mm (FWHM). The total intracranial volume (TIV) of each subject was calculated to use as a covariate for CV data.

For surface-based morphometry (SBM), we used the surface-preprocessing pipeline of the CAT 12 toolbox, which provides a fully automated method to estimate CT and the central surface of hemispheres based on the projection-based thickness method.^35^ This toolbox then allows the computation of multiple surface parameters, including CT, FD, and SD.^36^ These surface parameters of the left and right hemispheres were separately resampled and smoothed with a 15-mm FWHM Gaussian kernel. The software parcellated the cortex into 34 regions-of-interest (ROI) per hemisphere using Hammer’s atlas for CV data and the Desikan atlas for surface measures, such as CT, FD, and SD.^37, 38^ We then averaged the data extracted from each ROI in each participant for further analysis.

### Identification of an endophenotype pattern of cortical architecture

Since our aim was to identify a pattern of cortical architecture serving as an endophenotype from a given set of data, we formulated the problem in the following manner. Based on previous findings showing that individuals with ASD and their unaffected siblings share a pattern of alterations when compared to TD people,^23^ we recognized that the endophenotype was a measurable component satisfying the following conditions:^39^

1. Individuals with ASD have higher values than TD people (i.e., ASD > TD);
2. Unaffected siblings of individuals with ASD also have higher values than TD people (i.e., unaffected sibling > TD); and
3. TD sibling pairs have similar values (i.e., TD ⍰ TD sibling).

With respect to conditions (1) and (2), we could achieve our aim by conducting a classification analysis, in which cortical architectures capable of discriminating people with a high endophenotype value from people with a low endophenotype value were identified. Individuals with ASD and their unaffected siblings were regarded as having the endophenotype, while TD people and TD siblings were envisaged as not having the endophenotype throughout this manuscript.

Following our previous study,^39^ we used sparse logistic regression (SLR)^40^ as a multivariate classification approach. Briefly, SLR relies on a hierarchical Bayesian estimation, in which the prior distribution of each element of the parameter vector is represented as a Gaussian distribution. Based on the automatic relevance determination, irrelevant features are not used in the classification, since the respective Gaussian prior distributions have a sharp peak at zero. When implemented in SLR, this efficient feature-elimination method can mitigate the over-fitting problem derived from a small sample size with high-dimensional features. To evaluate the performance of the classifier, a leave-one-pair-out cross-validation (LOPOCV) was performed. Of note, a “pair” stands for sibling pairs. In each fold, all-but-one pair was used to train the SLR classifier, while the remaining pair was used for the evaluation. We controlled for age, FSIQ, and TIV for the CV data and age and FSIQ for the other cortical parameters.

To further examine the statistical significance of the classification accuracy, a permutation test was performed. While keeping the pair information, a permuted dataset was generated by shuffling the endophenotype label at each iteration. Then, we conducted LOPOCV to calculate the classification accuracy for the permuted dataset. This procedure was repeated 5,000 times to construct a null distribution. Statistical significance was set at *P* < 0.05.

### Binomial test

SLR selected a small number of relevant ROIs from a given set of data (see Results). To confirm that the ROIs selected by the classifier were not randomly selected, a binomial test examined the statistical significance of the selection counts. The classifier selected 8.93 ± 0.63 (mean ± standard deviation) out of 68 ROIs for each of the 30 validation folds (see Results). Thus, we assumed a binomial distribution *Bi* (*n, p*), where *n* stands for the number of validation folds (i.e., *n* = 30) and *P* stands for the probability of being selected from the set of ROIs (i.e., *P* = 9/68).

### Identification of neural correlates of a clinical ASD diagnosis

Once SLR identified a parameter that could capture an ASD endophenotype, we then performed a bootstrapping analysis to depict the neural correlates of a clinical ASD diagnosis among a given set of data. This analysis evaluates the difference in cortical 11 architecture between individuals with ASD and their unaffected siblings accounting for the distribution of the difference in SD between TD sibling pairs, following our previous study ^25^. First, we randomly assigned each of the TD sibling pairs into two groups and calculated the mean difference between the two groups, which provided the distribution of the difference in the SD of a given ROI between typical siblings. Because of the stringent threshold for statistical significance (see below), we repeated this procedure with 100,000 iterations. Then, we overlaid the mean difference in these measures between individuals with ASD and their unaffected brothers on the distribution of these measures between typical siblings. The statistical threshold for significance was set at either above 99.964 or below the 0.036 percentile, which was equivalent to *P* < 0.001 (=0.025/68) two-tailed.

### Relation between neural correlates of a clinical ASD diagnosis and clinical phenotype

To examine whether the brain parameters that emerged as neural correlates of a clinical ASD diagnosis were correlated with clinical symptoms, correlation analyses were performed. The analyses were performed with the mean SD of ROIs that showed significant results and subscores of the AQ among people with ASD (see Results).

## Results

### Identification of an ASD endophenotype

The SLR classifier with LOPOCV for the surface parameter of SD successfully dissociated participants with the endophenotype (i.e., individuals with ASD and their unaffected siblings) from the participants without an ASD endophenotype with a 73.3% 12 accuracy (sensitivity = 76.7% and specificity = 70.0%) and an area under the curve (AUC) of 0.75 (permutation test with 5,000 iterations, *P* < 0.001; Figure 1). In terms of the other parameters, specifically CV, CT, and FD, their classification accuracy and AUC were 48.3% (AUC = 0.53), 58.3% (AUC = 0.61), and 61.7% (AUC = 0.62), respectively. These results suggest that ASD endophenotype-related features are more evident for SD than for other surface-based parameters or CV. We therefore focused on the SD data in subsequent analyses.

**Figure 1.**
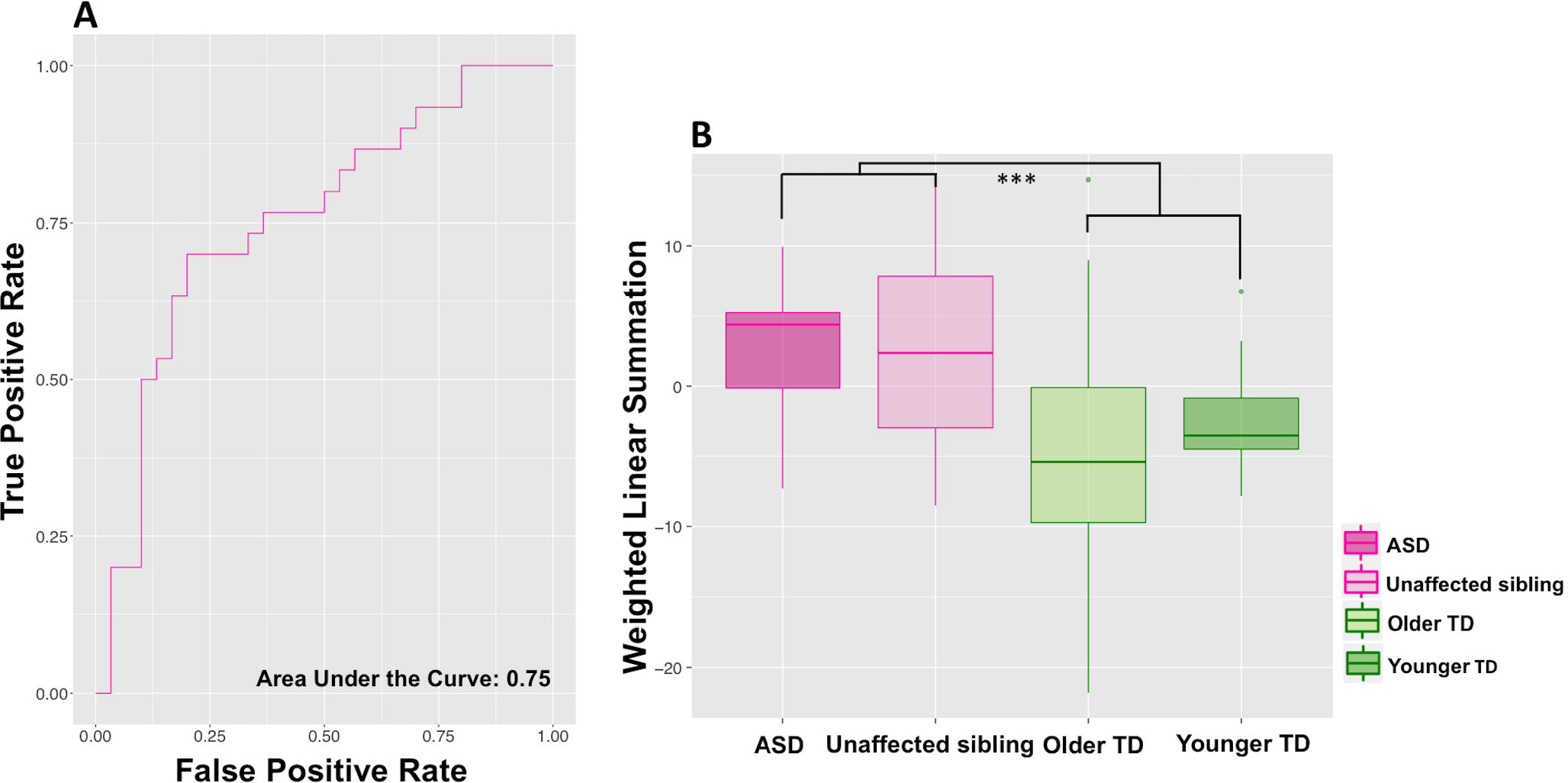
Classification results and post-hoc tests. (A) Receiver operating characteristic curve. A sparse logistic regression with leave-one-pair-out cross validation exhibited an area under the curve of 0.75. (B) Post-hoc paired t-tests demonstrated that there were no statistically significant differences between the individuals with autism spectrum disorder and their unaffected siblings in the weighted linear summation of selected sulcal depth (*t*-value = −0.15, *df* = 14, *P* = 0.89) as well as between typically developing pairs (*t*-value = −0.93, *df* = 14, *P* = 0.37). On the other hand, people with the endophenotype exhibited significantly higher values than those without the endophenotype (*t*-value = 3.7, *df* = 58, *P* < 0.001). ***: *P* < 0.001. Abbreviations: ASD: autism spectrum disorder, TD: typically developing.

We then performed post-hoc paired *t*-tests to examine whether the conditions (see Methods) were satisfied (Figure 1). A two-sample *t*-test showed statistically significant differences between participants with the endophenotype and those without (*t*-value = 3.7, *df* = 58, *P* < 0.001). In contrast, a paired *t*-test demonstrated that the WLS of the SD data selected in the classifier was not significantly different between individuals with ASD and their unaffected siblings (*t*-value = −0.15, *df* = 14, *P* = 0.89). The analysis also showed no significant difference between TD sibling pairs (*t*-value = −0.93, *df* = 14, *P* = 0.37). These results indicate that the WLS of the SD data selected by the classifier satisfies a set of conditions regarding endophenotype, but not clinical diagnosis.

### Binomial test for SD data

Furthermore, we investigated which SD data was stably selected by the SLR across LOPOCV. In each loop, the classifier selected 8.93 ± 0.63 out of 68 ROIs across 30 validation folds. We counted how many times each ROI was selected. Under the null hypothesis that nine brain regions were randomly selected from 68 ROIs, a binomial test was applied to examine the probability of the selection count. We found nine brain regions that were statistically significantly frequently selected (*P* < 0.05, Bonferroni corrected for 68 ROIs; Figure 2 and Table 3). For the nine ROIs, the selection count was 26.44 ± 2.54, while it was 0.50 ± 1.06 for the remaining ROIs. This result indicates that the nine ROIs were consistently selected across the validation folds.

**Figure 2.**
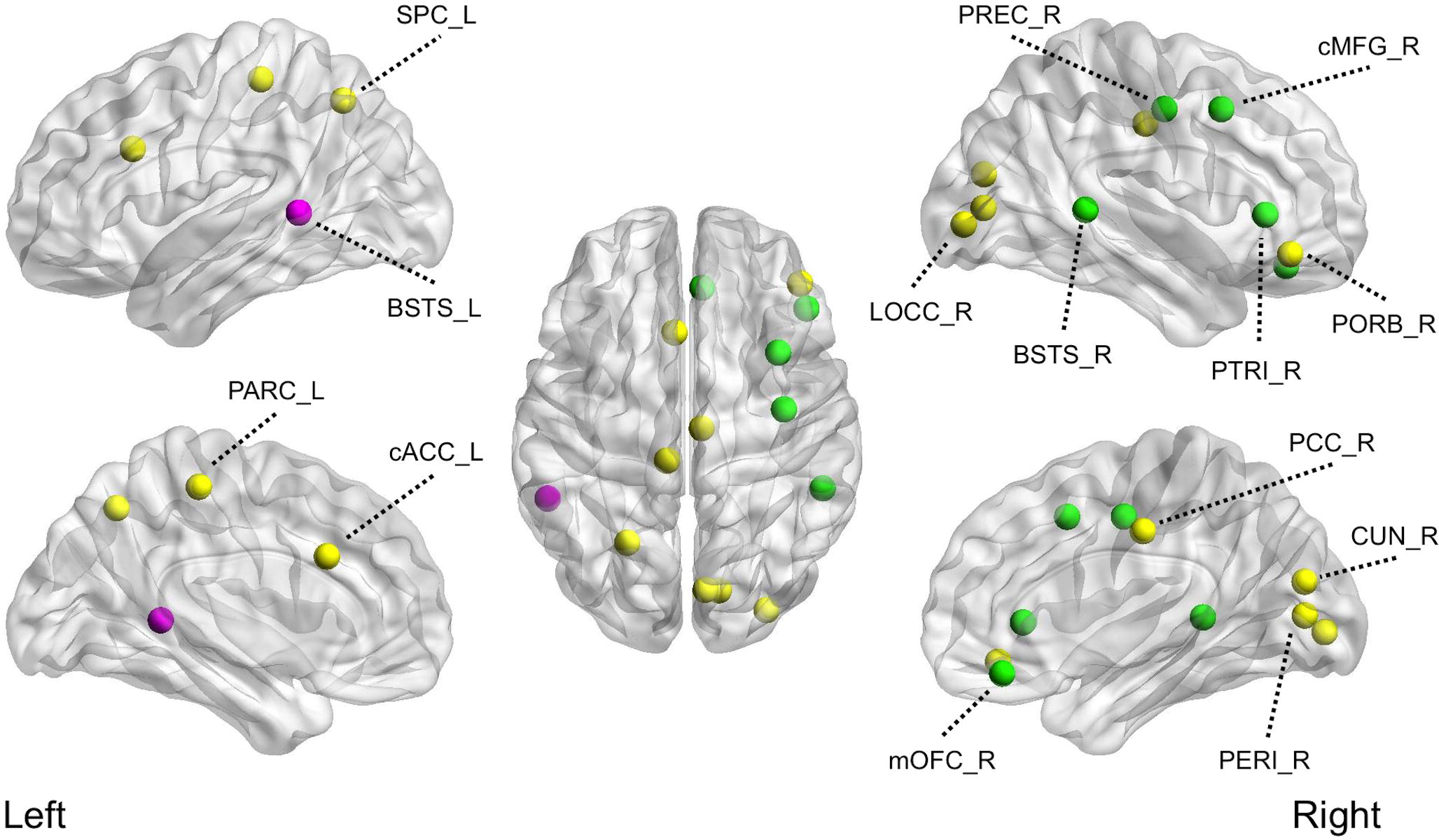
Results of multivariate machine-learning approach and bootstrapping analysis. The yellow circles represent the brain regions where the sulcal depth was associated with the ASD endophenotype. The green circles show the brain regions in which the neural correlates of an ASD diagnosis were associated with the sulcal depth. The pink circles represent a brain region associated with both the ASD endophenotype and neural correlates of an ASD diagnosis. Abbreviations: cACC_L, left caudal anterior cingulate cortex; CUN_R, right cuneus cortex; LOCC_R, right lateral occipital cortex; cMFG_R, right caudal middle frontal gyrus; mOFC_R, right medial orbitofrontal cortex; PARC_L, left paracentral lobule; PCC_R, right posterior cingulate cortex; PERI_R, right pericalcarine cortex; PORB_R; right pars orbitalis; PREC_R, right precentral gyrus; PTRI_R, right pars triangularis; SPC_L, left superior parietal cortex; pSTS_L, left posterior superior temporal sulcus; pSTS_R, right posterior superior temporal sulcus.

### Identification of neural correlates of a clinical ASD diagnosis

Focusing on SD, the analysis revealed that there were significantly large differences between individuals with ASD and their unaffected siblings, compared with those among TD sibling pairs, in six ROIs. The ROIs with significant differences included the left bank of superior temporal sulcus (<0.01 percentile), right pars triangularis, banks superior temporal sulcus, medial orbitofrontal cortex, precentral gyrus (<0.01 percentile) (see brain region details in Figure 2 and Table 3).

### Relationship between cortical measures correlated with a clinical ASD diagnosis and clinical phenotype

We examined the correlations between the SD values for the six ROIs that were significant in the bootstrapping analyses with five AQ subscales in the ASD group. The mean SD in the right bank of the superior temporal sulcus was correlated with the AQ attention switching/tolerance of change score (*r* = −0.655, *P* = 0.015). In addition, the mean SD in the left caudal middle frontal gyrus was correlated with the AQ social skills score (*r* = −0.562, *P* = 0.045).

## Discussion

The present study identified SD as a primary feature of cortical surface morphology that was representative of the endophenotype of ASD, compared with other cortical surface measures such as CV, CT, and FD. More specifically, a machine-learning approach for SD classified people with or without the ASD endophenotype with a 73.3% accuracy (sensitivity = 76.67% and specificity = 70.0%), while the classification accuracy of the other cortical parameters was relatively low. Furthermore, focusing on SD, a bootstrapping analysis successfully dissociated neural correlates of a clinical ASD diagnosis from an ASD endophenotype, which was not discriminable using the WLS of the SLR.

In this study, most of the ROIs identified as classification inputs were located within the “social brain”.^41, 42^ Although the unaffected siblings group was indistinguishable from the TD group at a clinical behavioral level, the predominance of the social brain in the endophenotype may reflect a predisposition to develop social communication deficits observed among people with ASD. Although atypical findings within the social brain are commonly observed,^4–6^ the better performance of SD over other parameters provides insight into the development of pathophysiology in the cortical architecture. Indeed, the cortical folding patterns are thought to reflect early patterns of cortico-cortical connectivity in the developing brain based on the theory of mechanical tension along long-distance axons.^43–45^ Therefore, we speculate that atypical cortical folding emerged as abnormal SD values among people with an increased genetic risk for developing ASD and may reflect alterations in connectivity involving these regions within the social brain during development.

CT represents dendritic arborization and pruning in gray matter in the brain^46^ and alterations in myelination at the merging of gray and white matter tissue.^47^ Regarding the developmental trajectories of CT in ASD, quadratic age trajectories composed of three phases have been proposed: accelerated expansion during early childhood, accelerated thinning during late childhood and adolescence, and lastly, decelerated thinning during early adulthood.^48, 49^ Notably, recent large sample studies have reported a dynamic CT pattern of group differences observed between children with ASD and those with TD over broad regions of the cortex, but with differences fading over adolescence to a virtually identical CT by young adult age.^9, 10^ Furthermore, a multivariate classification study using the Autism Brain Imaging Data Exchange dataset^50^ concluded that anatomical differences in cortical measures of CV, CT, and surface area offer very limited diagnostic value in ASD.^51^ When the samples are limited to only those obtained during adulthood, these prior lines of evidence may also support our finding that SD is potentially a predominant cortical surface parameter for capturing the ASD endophenotype.

The current study revealed a significantly large difference in SD between individuals with ASD and their unaffected siblings, compared with those among TD sibling pairs, within the “social brain.” Such area includes bilateral banks of the superior temporal sulcus (defined as the posterior aspect of the superior temporal sulcus; pSTS) and the right caudal middle frontal gyrus (MFG), the pars triangularis of the inferior frontal gyrus (IFG), and the medial orbitofrontal cortex (OFC). Interestingly, while atypical SD in the left pSTS was seen in both the endophenotype and neural correlates for a clinical diagnosis, atypical SD in the right pSTS was observed only in the clinical diagnosis. Furthermore, in the right pSTS, a negative correlation between the mean SD and the AQ attention switching/tolerance of change score was found, suggesting that a shallower pSTS may be associated with severe clinical symptoms. The pSTS (often the right pSTS) plays a central role in social cognition impairments among individuals with ASD, including biological motion perception, reading intentions from actions,^52, 53^ speech perception, audio-visual integration, perception of gaze and face processing.^54, 55^ By enrolling both siblings with and without an ASD endophenotype, the current study identified that atypical SD in the right pSTS may be a more specific feature representing an effect of the development of an ASD diagnosis, rather than a predisposition. These findings are consistent with a previous functional MRI study showing that atypical activity in the right pSTS serves as a biological marker of having ASD, rather than a genetic vulnerability to developing ASD.^56^

The results of the current study should be interpreted with caution. First, because of practical difficulties, we could not enroll monozygotic twins. Given that such individuals share genetic characteristics and share pre- and peri-natal factors as well as a similar environment after birth, monozygotic twins would fit the aim of the current project more than full siblings. In addition, the number of participants per group was relatively small, although we obtained data from 60 participants in total. The relatively small number of participants might have induced overfitting, while the SLR might have mitigate it. Future research with a larger sample size is expected to overcome these limitations.

In summary, atypical cortical folding, represented by SD, within the social brain was identified as a core feature associated with the endophenotype of ASD. Using SD, we successfully dissociated the neural correlates of the development of a clinical ASD diagnosis from its endophenotype, recruiting not only siblings discordant for ASD, but also pairs of typically developing siblings.

## Acknowledgements

The authors thank Kei Kamiya and Tomomi Kikuya for their contributions to the data collection. This work was partially supported by JSPS KAKENHI [grant number JP 25861030] and the Joint Usage/Research Program of the Medical Institute of Developmental Disabilities Research, Showa University (BY). This work was also partially supported by the Brain Mapping by Integrated Neurotechnologies for Disease Studies (Brain/MINDS) from the Japan Agency for Medical Research and Development (AMED) (RH).

## Author contributions

BY, IT, and YA designed the study, analyzed data, and wrote the paper. BY and IT collected data. All the other authors provided critical revision of the paper.

