## Supplementary material for "Cortical surface architecture endophenotype and correlates of clinical diagnosis of autism spectrum disorder": Table3

**Table 3.** Brain regions of sulcal depth associated with an ASD endophenotype and neural correlates of a clinical ASD diagnosis

| **ASD endophenotype** |  |  | **Neural correlates of a clinical ASD diagnosis** |  |
| --- | --- | --- | --- | --- |
| Brain region (multivariate classification approach) | Mean weight |  | Brain region (Bootstrapping analysis) | Percentile |
| Left bank superior temporal sulcus | -6.4 |  | Left bank superior temporal sulcus | <0.01 |
| Left caudal anterior cingulate cortex | -5.8 |  |  |  |
| Left paracentral lobule | 3.1 |  |  |  |
| Left superior parietal cortex | -5.0 |  |  |  |
| Right cuneus | 4.9 |  |  |  |
| Right lateral occipital cortex | -7.6 |  |  |  |
| Right pars orbitalis | -5.2 |  |  |  |
| Right pericalcarine cortex | -4.5 |  |  |  |
| Right posterior cingulate cortex | 4.4 |  |  |  |
|  |  |  | Right bank superior temporal sulcus | <0.01 |
|  |  |  | Right caudal middle frontal gyrus | 0.01 |
|  |  |  | Right pars triangularis | <0.01 |
|  |  |  | Right medial orbitofrontal cortex | <0.01 |
|  |  |  | Right precentral gyrus | <0.01 |

Abbreviations: ASD: autism spectrum disorder
