## Supplementary material for "Cortical surface architecture endophenotype and correlates of clinical diagnosis of autism spectrum disorder": Table2

| **Table 2.** ASD symptoms of people with an ASD endophenotype | | | | | | | |
| --- | --- | --- | --- | --- | --- | --- | --- |
|  | Individuals with ASD | | Unaffected siblings | | *Statistics (a)* | | |
|  | (n = 15) | | (n = 15) | |  |  |  |
|  | Mean | SD | Mean | SD | df | *t* value | *P* value |
| AQ |  |  |  |  |  |  |  |
| AS | 7.4 | 1.6 | 3.9 | 1.8 | 12 | 6.3 | <0.001 |
| ATD | 5.4 | 1.9 | 3.2 | 1.9 | 12 | 2.67 | 0.02 |
| COM | 7.6 | 1.6 | 3.6 | 2.6 | 12 | 4.16 | 0.001 |
| IMG | 6 | 2 | 4.1 | 1.4 | 12 | 2.31 | 0.04 |
| SS | 7.9 | 2 | 4.5 | 2.7 | 12 | 4.26 | 0.001 |
| Total | 34.2 | 6.2 | 19.4 | 6.9 | 12 | 6.27 | <0.001 |
| ADI-R* |  |  |  |  |  |  |  |
| Social interaction | 20.7 | 6.3 | 0.8 | 1.4 | 13 | 11.56 | <0.001 |
| Communication | 12.6 | 5 | 0.4 | 1.1 | 13 | 8.83 | <0.001 |
| RRB | 3.9 | 2 | 0 | 0 | 13 | 7.02 | <0.001 |
| ADOS** |  |  |  |  |  |  |  |
| Communication | 4.45 | 1.13 | - | - | - | - | - |
| Social interaction | 7.91 | 2.12 | - | - | - | - | - |
| Communication + Social interaction | 12.36 | 2.73 | - | - | - | - | - |
| RRB | 0.18 | 0.4 | - | - | - | - | - |
| (a) The statistics show the results of comparisons between adult males with ASD and their unaffected brothers. Abbreviations: ADOS-2: Autism Diagnostic Observation Schedule Second Edition, ADI-R: Autism Diagnostic Interview-Revised, AQ: autism-spectrum quotient, AS: attention switching/tolerance of change, ASD: autism spectrum disorder, ATD: attention to detail, COM: communication skills, IMG: imagination, IQ: intelligence quotient, RRB: restricted repetitive behaviors, SD: standard deviation, SES: socioeconomic status, SS: social skills, TD: typical development. *The ADI-R score was missing for one person. **ADOS scores are missing for four people. | | | | | | | |
