## Supplementary material for "Cortical surface architecture endophenotype and correlates of clinical diagnosis of autism spectrum disorder": Table1

| **Table 1.** Participant characteristics | | | | | | | | | | | | | | | |
| --- | --- | --- | --- | --- | --- | --- | --- | --- | --- | --- | --- | --- | --- | --- | --- |
|  | People with endophenotype (n = 30) | | | | Statistics (a) | | |  | People without endophenotype (n = 30) | | | | Statistics (d) | | |
|  | Individuals with ASD | | Unaffected siblings | | df | *t* value | *P* value |  | TD (b) | | TD (c) | | df | *t* value | *P* value |
|  | (n = 15) | | (n = 15) | |  |  |  |  | (n = 15) | | (n = 15) | |  |  |  |
|  | Mean | SD | Mean | SD |  |  |  |  | Mean | SD | Mean | SD |  |  |  |
| Age (years) | 28.3 | 6.1 | 28 | 7.3 | 14 | 0.45 | 0.657 |  | 28.4 | 6.5 | 25.1 | 5.3 | 14 | 5.62 | <0.001 |
| Handedness | 61.8 | 68.4 | 99.3 | 2.9 | 14 | 2.11 | 0.054 |  | 89.2 | 26.4 | 80.7 | 51.4 | 14 | 0.55 | 0.594 |
| Full IQ | 111.8 | 16 | 107.4 | 13.2 | 14 | 0.77 | 0.452 |  | 115.9 | 15.7 | 114.9 | 12.1 | 14 | 0.24 | 0.815 |
| Verbal IQ | 106.3 | 31.2 | 107.1 | 16.8 | 14 | 0.08 | 0.94 |  | 117.1 | 15.9 | 116.7 | 12.5 | 14 | 0.18 | 0.859 |
| Performance IQ | 108.1 | 16.4 | 105.3 | 9.8 | 14 | 0.62 | 0.547 |  | 109.4 | 13.6 | 107.7 | 7.9 | 14 | 0.42 | 0.678 |
| SES (e) | 5.5 | 1.1 | 5.5 | 1.1 | 14 | 0.2 | 0.843 |  | 5.9 | 1.2 | 5.7 | 1.1 | 13 | 0.46 | 0.655 |
| (a) The statistics show the results of comparisons between adult males with ASD and their unaffected brothers. (b) Older siblings. (c) Younger siblings. (d) The statistics show the results of comparisons between typical older and younger siblings. (e) A higher score indicates a lower socioeconomic status.^57^ Abbreviations: ASD: autism spectrum disorder, IQ: intelligence quotient, SD: standard deviation, SES: socioeconomic status, TD: typical development. | | | | | | | | | | | | | | | |
